# Robust Variation in Infant Gut Microbiome Assembly Across a Spectrum of Lifestyles

**DOI:** 10.1101/2022.03.30.486467

**Authors:** Matthew R. Olm, Dylan Dahan, Matthew M. Carter, Bryan D. Merrill, Brian Yu, Sunit Jain, Xian Dong Meng, Surya Tripathi, Hannah Wastyk, Norma Neff, Susan Holmes, Erica D. Sonnenburg, Aashish R. Jha, Justin L. Sonnenburg

## Abstract

Infant microbiome assembly is intensely studied in infants from industrialized nations, but little is known about this process in populations living non-industrialized lifestyles. In this study we deeply sequenced infant stool samples from the Hadza hunter-gatherers of Tanzania and analyzed them in a global meta-analysis. Infant microbiomes develop along lifestyle-associated trajectories, with over twenty percent of genomes detected in the Hadza infant gut representing phylogenetically diverse novel species. Industrialized infants, even those who are breastfed, have microbiomes characterized by a paucity of *Bifidobacterium infantis* and gene cassettes involved in human milk utilization. Strains within lifestyle-associated taxonomic groups are shared between mother-infant dyads, consistent with early-life inheritance of lifestyle-shaped microbiomes. The population-specific differences in infant microbiome composition and function underscore the importance of studying microbiomes from people outside of wealthy, industrialized nations.

**Recognition of work on indigenous communities:** Research involving indigenous communities is needed for a variety of reasons including to ensure that scientific discoveries and understanding appropriately represent all populations and do not only benefit those living in industrialized nations. Special considerations must be made to ensure that this research is conducted ethically and in a non-exploitative manner. In this study we performed deep metagenomic sequencing on fecal samples that were collected from Hadza hunter-gatherers in 2013/2014 and were analyzed in previous publications using different methods (*1, 2*). A material transfer agreement with the National Institute for Medical Research in Tanzania ensures that stool samples collected are used solely for academic purposes, permission for the study was obtained from the National Institute of Medical Research (MR/53i 100/83, NIMR/HQ/R.8a/Vol.IX/1542) and the Tanzania Commission for Science and Technology, and verbal consent was obtained from the Hadza after the study’s intent and scope was described with the help of a translator. The publications that first described these samples included several scientists and Tanzanian field-guides as co-authors for the critical roles they played in sample collection, but as no new samples were collected in this study, only scientists who contributed to the analyses described here were included as co-authors in this publication. It is currently not possible for us to travel to Tanzania and present our results to the Hadza people, however we intend to do so once the conditions of the COVID-19 pandemic allow it.

## Main Text

The human gut microbiome undergoes a complex process of assembly beginning immediately after birth (*3*). New microbes attempting to engraft within this community often depend upon niches established by previous colonizing species, and thus the final adult microbiome composition may be contingent upon the species acquired early in life. The microbiome assembly process of infants living in industrialized nations is well-studied, and tends to follow a series of characterized steps that lead to the low-diversity gut microbiome composition characteristic of industrialized adults (*4*). The microbiome assembly process that occurs in infants living non-industrialized lifestyles (which results in the characteristically diverse adult microbiomes of non-industrialized adults (*2*)) is largely unknown (*5*). Of particular interest are the timing at which the microbiomes of infants from different lifestyles diverge, the microbes and functions that are characteristic of infants from different lifestyles, and whether there are differences in the taxa that are vertically transmitted from mothers to infants and seed the microbiome assembly process.

To address these questions we performed metagenomic sequencing on infant fecal samples from the Hadza, a group of modern hunter-gatherers in Sub-Saharan Africa (*1, 6*). The Hadza inhabit semi-nomadic bush camps of approximately 5 to 30 people and exhibit a moderate level of community child rearing within the camps (*7*). Hadza infants are breastfed early in life and then weaned onto a diet that consists of baobab powder, animal fat, and pre-masticated meat at approximately 2–3 years old (*8, 9*). In this study we performed i) a global meta-analysis of infant microbiome samples sequenced using 16S rRNA amplicon sequencing, including 62 Hadza infant fecal samples, in order to contextualize the Hadza infant microbiome with as many samples as possible, and ii) deep metagenomic sequencing on 39 Hadza infant fecal samples in order to assess sub-species variation, functional potential, and patterns of vertical transmission **(Table S1, S2)**.

To assess inter-individual variation of infants across lifestyles, we curated a global dataset of 16S rRNA sequencing samples from 1,900 healthy infants aged 0-36 months from 18 populations (including the Hadza samples described above) (*1, 2, 4, 10*–*14*) (**Fig. S1, S2**). Populations were categorized as practicing industrialized, transitional, or non-industrialized lifestyles using the Human Development Index (HDI) and other lifestyle characteristics (*15, 16*); see methods for details. We created a UniFrac-based ordination from all 1,900 samples **(Fig. 1A)** and found that the main ordination axis of variation is strongly associated with age (EnvFit; n=1900; R^2^ = 0.43; P = 0.001) and the second axis of variation is strongly associated with lifestyle (EnvFit; n=1900; R^2^ = 0.50; P = 0.001). DNA extraction methods, or other study-specific aspects of data generation, may contribute to some of the differences in data between studies. Comparison of two populations within the same country but with different lifestyles (the Bassa from Nigeria (non-industrialized lifestyle) and city dwelling Nigerian infants (transitional lifestyle)) demonstrate that shared lifestyle affects microbiota composition more than geographic proximity (**Fig. 1A**, right panel**; Fig. S3**). Similarly, the Tsimane infants in Bolivia harbor a microbiota more similar to Hadza and Bassa infants than to infants from other lifestyles in South America (**Fig. S3**). The microbiome of infants living industrialized lifestyles diverges from those living transitional and non-industrialized within the first 6 months of life, and the microbiomes of infants living transitional versus non-industrialized lifestyles diverge at ∼30 months of life **(Fig. 1B)**. A lack of complete metadata precluded us from testing whether this is due to differences in feeding practices between lifestyles. Among infants living transitional lifestyles, intermediate trajectories are exhibited by populations on the boundaries of industrialized or non-industrialized lifestyles (**Fig. 1B**, dashed lines), highlighting the apparent sensitivity of infant microbiota development to lifestyle-related factors.

**Fig. 1.**
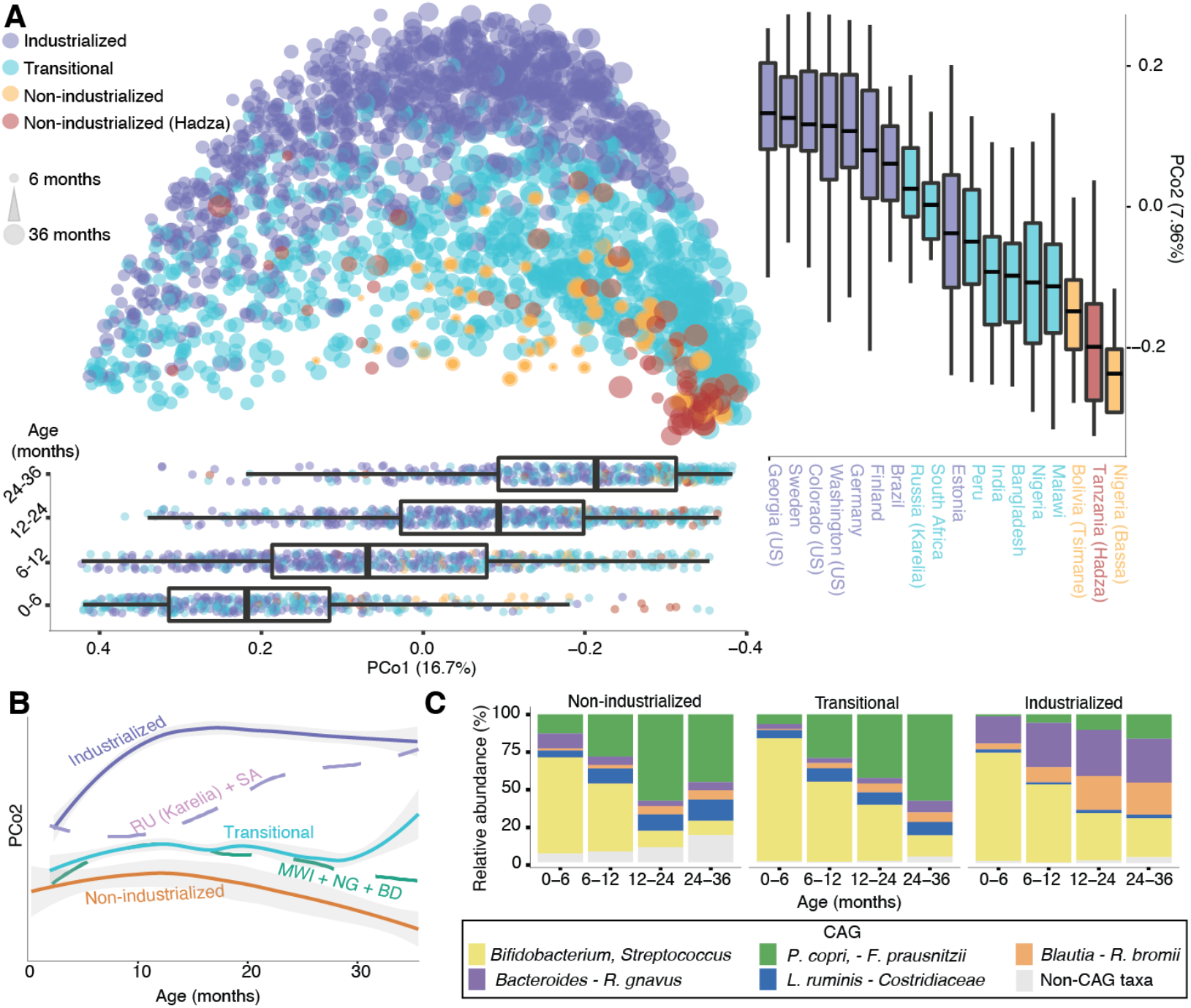
Age and lifestyle are associated with infant microbiome composition. **(A)** Unweighted UniFrac dissimilarity PCoA (top left panel) of 1900 infant fecal samples across 18 populations based on amplicon sequence variant (ASV) abundance. Point color indicates lifestyle, point size is proportional to age in months. Boxplots show the distribution of indicated age groups along PCo1 (bottom) and cohorts along PCo2 (right). **(B)** PCo2 versus sample age for the three lifestyle categories (solid lines) and specific indicated subpopulations (dashed lines). Upper purple dashed line includes Russia (Karelia) and South Africa (RU (Karelia)+SA) and the lower green dashed line includes Bangladesh, Malawi, and Nigeria (Urban) (MWI+NG+BD). The middle transitional line contains all transitional samples. Lines are the smoothed conditional mean of PCo2 loadings (loess fit). **(C)** Fractional abundance of co-abundance groups (CAGs) by age group and lifestyle. Taxa in annotation are the most abundant taxa in a CAG.

Members of the gut microbiome are often metabolically or ecologically linked, for example in the respective production and consumption of metabolites. We identified five microbial co-abundance groups (CAGs) in our dataset using a network inference method (*17, 18*), which together account for an average of 93.8% of the microbiota composition per sample (**Fig. 1C; Fig. S4**). The *Bifidobacterium-Streptococcus* CAG dominates infants from all lifestyles in early life (0-6 months) **(Fig. 1C)** and yields over time in a lifestyle-specific manner. A *Bacteroides-Ruminocccocus gnavus* CAG is enriched in industrialized infants whereas a *Prevotella-Faecalibacterium* CAG is enriched in infants living transitional / non-industrialized lifestyles (**Fig. 1C**). These differences in dominant CAGs by lifestyle become more pronounced over time and mirror taxonomic tradeoffs observed in late infancy (*19*) and compositional differences found in adult microbiomes (*1*).

We next used our deep metagenomic sequencing data of Hadza infant fecal samples, in comparison to other available infant metagenomes, to assess microbiome-encoded functional differences across samples (see methods for details). Broad lifestyle-associated differences were seen in the overall functional capacity of the infant microbiomes, along with age-related microbiome development **(Fig. 2A)**. These metagenomic data are consistent with 16S rRNA amplicon-based analysis **(Fig. 1A)** illustrating that these phylogenetic differences have functional consequences.

**Fig. 2.**
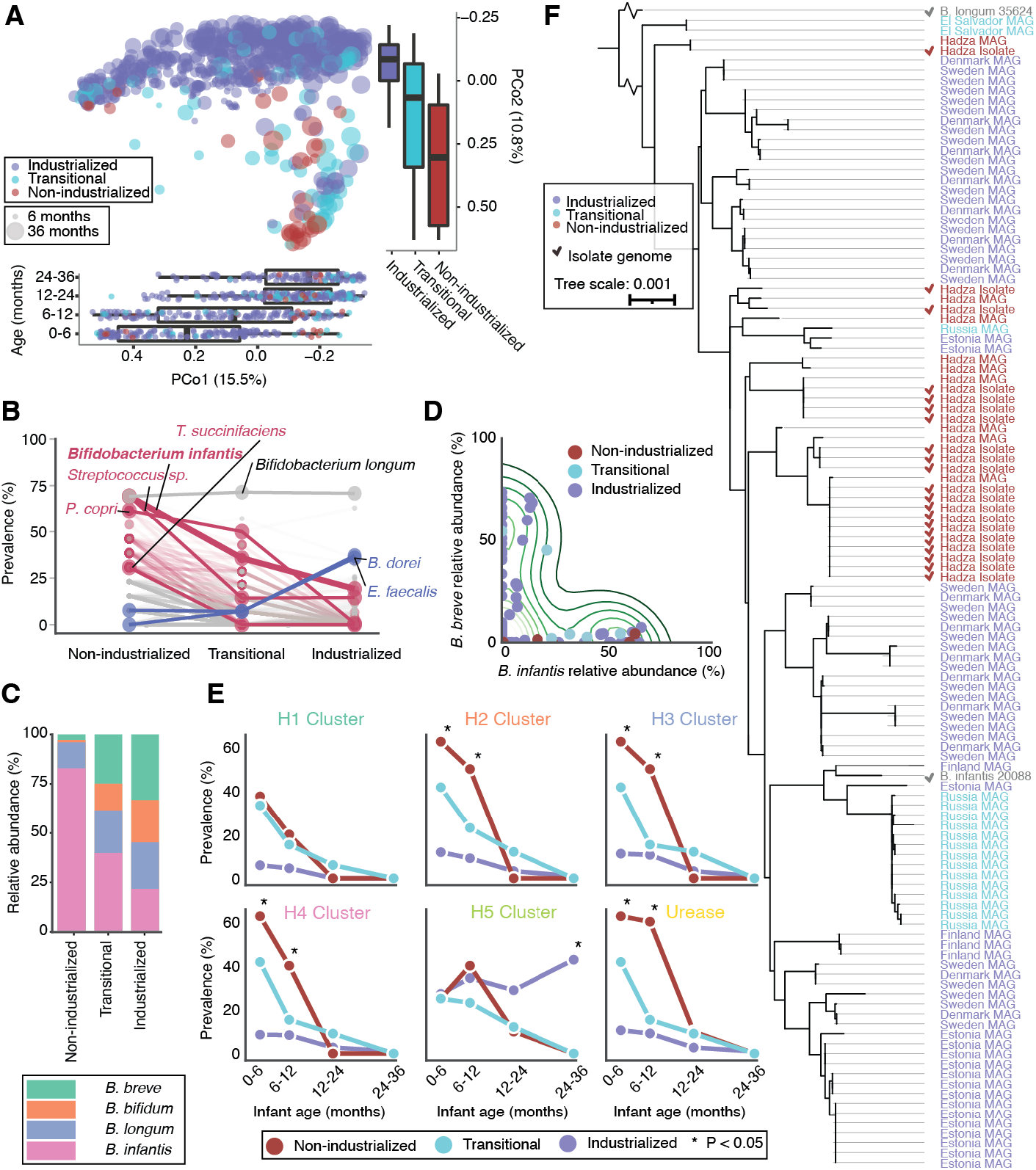
Age and lifestyle are associated with infant microbiome functions. **(A)** PCoA based on 682 infant fecal metagenomes described at the gene abundance level in RPKM. Points are colored by lifestyle. Size indicates infant age in months. Boxplots (bottom panel) show the distribution of indicated age groups in months along PCo1.Boxplots (right panel) show the distribution of each lifestyle along PCo2. The main axis of variation in this gene-based ordination is significantly associated with age (EnvFit; R2 = 0.30; n=679; P = 0.001) and the second axis of variation is significantly associated with lifestyle (EnvFit; R2=0.35; n=679; P=0.001). **(B)** Fractional prevalence of species across lifestyles among 0-6 month old infants. Select VANISH (red) and BloSSUM (blue) (those with lowest adj-P) species are highlighted. *B. infantis* is shown in bold. “Other” (gray) taxa are those that are not significantly different by lifestyle. **(C)** Relative representation of four common *Bifidobacterium* species in 0-6 month olds by lifestyle. **(D)** Scatterplot of *B infantis* versus *B. breve* abundance among 0-6 month old infants. Contour lines display the kernel density estimation (KDE). **(E)** Prevalence of HMO-utilization clusters across ages and lifestyles. Clusters are considered present if all genes in the cluster are detected above a variable coverage threshold (to ensure that results are robust to differences in sequencing depth; see methods for details). * = adj-P < 0.05; Fisher’s exact test with false discovery rate correction; non-industrialized versus industrialized. **(F)** Phylogenetic tree of *B. infantis* genomes based on universal single copy genes. Genome names are colored based on lifestyle of origin. Isolate genomes are marked with a checkmark. Public reference genomes for *B. longum* and *B. infantis* are included in gray text.

Hadza infant metagenomes were assembled and binned into metagenome-assembled genomes (MAGs) to investigate taxonomic novelty and sub-species variation. Of the 745 microbial species assembled from Hadza infant metagenomes, 23% (n=175) represent novel species according to UHGG **(Table S7**). These novel species belong to phylogenetically diverse taxonomic groups **(Fig. S5A)**, 88.6% (n=155) were recovered from multiple Hadza samples **(Fig. S5B)**, and their genome quality is similar to genomes in UHGG **(Fig. S5C)**. To assess prevalence via read mapping, MAGs were integrated with genomes recovered from Hadza adults (*20*) and public genomes from the human gut (*21*) into a comprehensive database of 5,755 species-representative genomes (see methods for details). Overall, 23.4% of microbial species detected in the Hadza infants represent novel species **(Table S3)**. These data support that, like the adult Hadza gut, the Hadza infant gut contains extensive previously-uncharacterized diversity.

The taxonomic specificity afforded by metagenomic sequencing allowed us to identify particular species that are depleted or enriched in infants living industrialized versus non-industrialized lifestyles. Microbial species exhibiting these patterns are termed VANISH (volatile and/or negatively associated in industrialized societies of humans) and BloSSUM (bloom or selected in societies of urbanization/modernization) species, respectively (*2*). 310 VANISH and 12 BloSSUM species were identified among the infants in this analysis (**Table S4, Fig. S6)** (see methods for classification details). 63 VANISH species are effectively extinct (i.e., never detected) in industrialized and transitional lifestyle infants. Two asymmetries are apparent among VANISH and BloSSUM species across lifestyles. First, VANISH species are more numerous and abundant than BloSSUM taxa. VANISH species collectively comprise on average 36.6% (± 2.16%) of non-industrialized lifestyle infant microbiomes by relative abundance throughout infancy, while BloSSUM species comprise 7.73% (± 0.49%) in industrialized lifestyle infants (P=8.2e-21; n=569 industrialized infants; n=39 non-industrial infants; Wilcoxon rank-sum test) (**Fig. S7**). Second, VANISH species are over-represented shortly after birth whereas BloSSUM species occur primarily later in infancy. Many VANISH species (45.2%; 140/310) are present at 0-6 months in non-industrialized infants, while few BloSSUM species are detected this early in industrialized lifestyle infants (16.7%; 2/12) (**Fig. 2B**). Together these patterns suggest that more species disappear than appear as lifestyles become more industrialized, and that differences in the very early life microbiome (0-6 months) may engender alternative trajectories of microbiome assembly.

Amplicon and metagenomic data both show that *Bifidobacterium* is the most prevalent taxon in early life (**Fig. 1C, Fig. 2B**), prompting us to next investigate variation in *Bifidobacterium* species and strains across lifestyles. In the first 6 months of life we found that non-industrial lifestyle infants are dominated by *Bifidobacterium infantis* (also known as *Bifidobacterium longum* subsp. *infantis*) **(Fig. 2C)**, a prolific utilizer of human milk oligosaccharides (HMOs), associated with human health, and commonly used in probiotic supplements (*22*). Infants living transitional and industrialized lifestyles have expanded representation of other *Bifidobacterium* species with limited ability to degrade HMOs (**Fig. 2C**). At 0-6 months of life, *B. infantis* is significantly decreased in industrial microbiomes (**Fig. 2C;** adj-P = 0.04; n=151 industrialized infants; n=27 non-industrial infants; Wilcoxon-ranked) and its abundance in transitional 0-6 month infants is at an intermediate state between non-industrial and industrialized infants. *Bifidobacterium breve*, a species capable of limited HMO degradation (*23*), is the most abundant *Bifidobacterium* species in industrialized infants. Interestingly, *Bifidobacterium infantis* is anti-associated with *Bifidobacterium breve* in infants across lifestyles **(Fig. 2D;** correlation=-0.46, P=4.1E-5, n=73 infants, spearman two-sided hypothesis test). Even industrialized infants that have *Bifidobacterium infantis* have low levels of *Bifidobacterium breve* **(Fig. 2D**; correlation=-0.41, P=1.0E-3, n=62 industrialized infants, spearman two-sided hypothesis test), suggestive of competitive exclusion.

To determine whether these species-level differences result in community-wide differences in HMO degradation capacity, we mapped our metagenomic reads to the most well-characterized genetic clusters for human milk utilization (**Table S5**). Five of these clusters are involved in HMO degradation (H1 - H5) and one cluster is involved in nitrogen scavenging (referred to as the “urease” cluster) (*22, 24*), and recent studies have linked their expression in the infant gut microbiome with systemic immunological health outcomes (*25*). Five of the six clusters were more prevalent in non-industrialized than industrialized infants, and their prevalence among transitional infants was in between these two extremes, in all cases showing a pattern of decreasing representation as infants age **(Fig. 2E)**. The H5 cluster, however, is found at the same prevalence level in infants from all lifestyles in early life, but exhibits continued persistence only in infants from industrialized lifestyles **(Fig. 2E)**. The H5 cluster encodes an ABC-type transporter known to bind core HMO structures, and we found this cluster in 119 of 129 *B*.*breve* MAGs and 41of 69 *B. infantis* MAGs recovered from industrialized infants (P = 1.4E-9, Fisher’s exact test). The persistence of the H5 cluster beyond 12 months in industrialized infants, a time period when breastfeeding is less common in these populations, suggests this cassette of genes exists in genomes that are not reliant upon breastfeeding. Breast milk consumption among industrialized infants reduces, but does not eliminate, lifestyle-associated differences in *Bifidobacterium infantis* and HMO-degradation cassette prevalence **(Fig. S8)**.

Beyond the species-level *Bifidobacterium* differences described above, we next leveraged the assembly-based metagenomic analysis performed in this study to investigate strain-level differences among *B. infantis* genomes recovered from infants living different lifestyles. *B. infantis* MAGs from infants aged 0-1 years old (n=96 MAGs) have several functional differences, including i) enrichment in non-industrialized versus industrialized infants of Glycoside Hydrolase Family 163 (GH_163), a CAZyme involved in the utilization of complex N-glycans (including those found on immunoglobulins) (*26*) **(Fig. S9A)**, ii) three Pfams that differ in prevalence in infants from different lifestyles **(Fig. S9C)**, and iii) the identification of four gene clusters at higher prevalence in *B. infantis* genomes found in the Hadza versus industrialized infants **(Fig. S9D)**. To verify the metagenome-based findings, we isolated and sequenced 20 *B. infantis* strains from the same Hadza infant fecal samples (**Table S7**). GH_163 and all four gene clusters showed enrichment among Hadza *B. infantis* isolates as compared to public reference genomes **(Fig. S9)**, consistent with our MAG-based findings. Finally, strong lifestyle-specific phylogenetic clustering was observed among *B. infantis* isolate sequences and MAGs **(Fig. 2F)**. This observation of strong region-specific phylogenetic signals **(Fig. 2F)** could be a result of long-term, multi-generational vertical transmission (*27*).

To assess the extent of vertical strain transmission among the Hadza, we deeply sequenced fecal samples from Hadza mothers of infants in this study (n=23 Hadza dyads). Detailed strain-tracking analysis was performed using inStrain (*28*) with a threshold for identical strains of 99.999% popANI (**Table S6**). Dyad pairs share far more strains on average than non-dyad pairs (6.4 vs 0.3 strains, respectively), and a higher percentage of infant strains are shared between an infant and their respective mother versus with another mother (12.4% vs 0.5%, respectively) (*p* < 0.001, Wilcoxon rank-sum test) **(Fig. 3A)**. Remarkably, Hadza non-dyads living in the same bush camp also share more strains and a higher proportion of strains than those living in different bush camps **(Fig. 3A)** (*p* < 0.001, Wilcoxon rank-sum test), consistent with previously-reported increased rates of strain sharing within Fijian social networks. (*29*). Vertically shared strains were detected among a range of different phyla in the Hadza, with Bacteroidota and Cyanobacteria having more vertically shared strains than other phyla, and Firmicutes having less **(Fig. 3B**; Fisher’s exact test with false discovery rate correction). Industrialized infants also exhibit significantly increased and decreased vertical strain sharing of Bacteroidetes and Firmicutes, respectively (*30*). Together these results suggest that community interaction during rearing of infants and bush camp microenvironments such as water source may play important roles in increasing transmission of strains to infants, and is consistent with proximity and social interactions propagating group-microbial sharing (*31*).

**Fig. 3.**
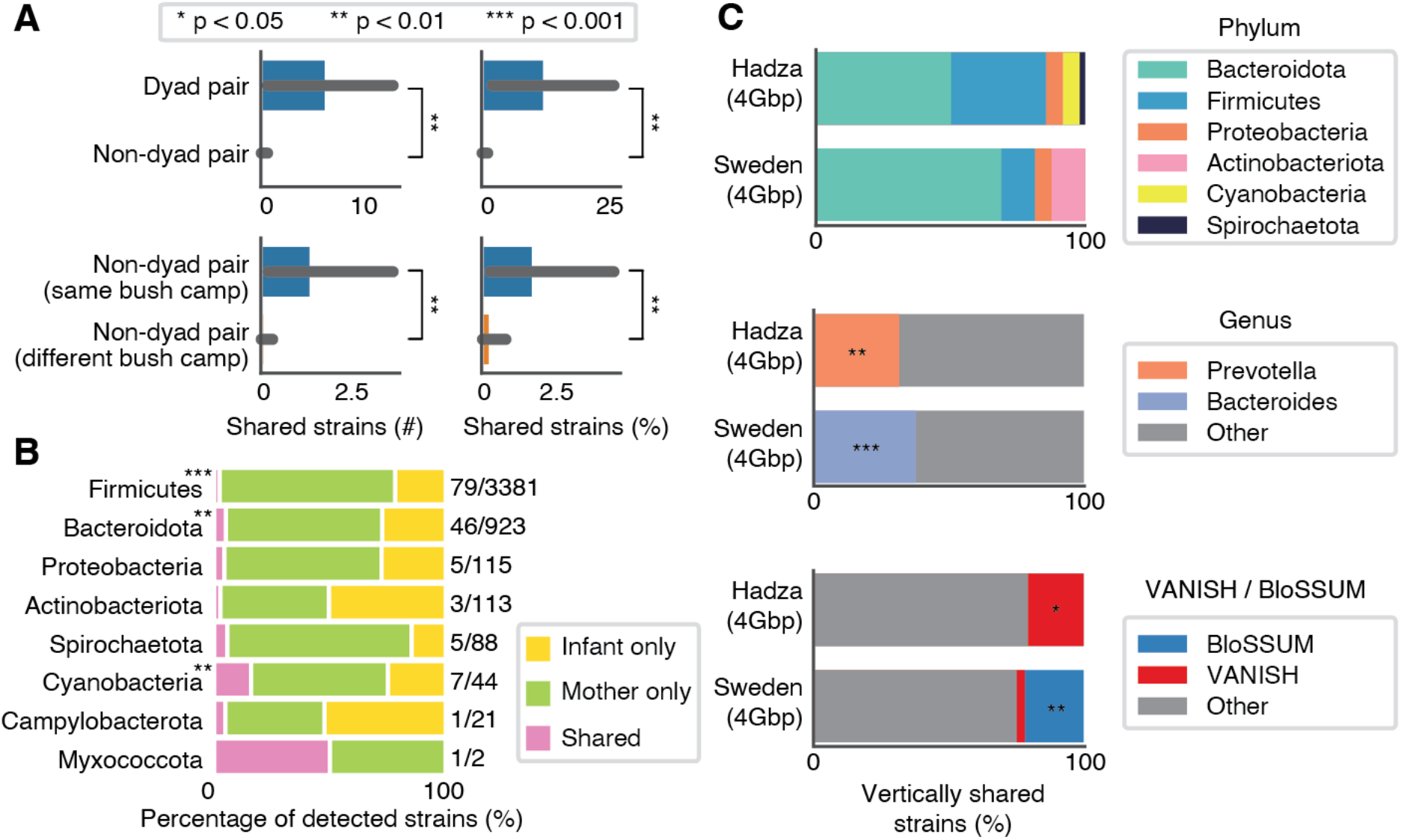
Strain sharing between mother-infant dyads and non-dyads is lifestyle-specific. **(A)** The mean strains shared (left) and the percentage of infant strains found in mothers (right) in mother-infant dyads versus mother-infant non-dyads (top) and non-dyads from the same bushcamp versus non-dyads from different bushcamps (bottom). Error bars represent standard error (* = adj-P < 0.05; ** = adj-P < 0.01; *** = adj-P < 0.001; Wilcoxen rank-sums test). **(B)** The percentage of strains detected in all Hadza mothers and infants and whether they are detected in infants only, mothers only, or shared within a mother / infant dyad (“Shared”) categorized by phylum. Numbers to the right of bars indicate the number of vertically shared strains over the number of strains detected in either infant or maternal samples. Phyla with a significant difference in the percentage of vertically transmitted strains as compared to all other phyla are marked with asterisks (Fisher’s exact test with p-value correction). **(C)** Percentage of vertically transmitted strains in Hadza and Swedish cohorts by phylum (top), genus (middle; only genera with significant differences shown), and VANISH / BloSSUM (bottom). All metagenomes were subset to 4Gbp for this analysis to reduce any biases associated with sequencing depth. Taxa that are significantly enriched in either cohort are marked with an asterisk (* = adj-P < 0.05; ** = adj-P < 0.01; *** = adj-P < 0.001; Fisher’s exact test).

To address whether lifestyle-dependent divergence of infant microbiotas could be explained by strain sharing between mothers and their infants, we performed the same detailed strain-tracking analysis on a previously sequenced dataset of 100 mother-infant dyads from Sweden (*32*). Swedish infants born via C-section were excluded from this analysis (n=17 eliminated; n=83 included). Infants in Swedish and Hadza dyads had average ages of 1.01 ± 0.00 and 0.95 ± 0.21 years old, respectively (P=0.04, Wilcoxon rank-sum test); one difference between these studies is that Swedish mothers were sampled immediately after birth while Hadza mothers were sampled contemporaneously with infants. To identify differences between strains shared among Hadza versus Swedish dyad pairs, we performed *in silico* rarefaction to account for differences in sequencing depth between the studies. Bacteria of the phylum Bacteroidota (also known as Bacteroidetes) were the most commonly vertically transmitted strains in both populations **(Fig. 3C)**. VANISH bacteria and bacteria of the genus Prevotella made up a higher portion of vertically shared strains in the Hadza, while bacteria of the genus Bacteroides and BloSSUM taxa were more commonly shared among Swedish dyads (Fisher’s exact test; P < 0.01) **(Fig. 3C)**. While we cannot exclude the possibility that the small difference in infant age between populations plays a role, the differences seen in infants largely mirror lifestyle-related compositional differences observed among adults, consistent with the finding that species that were more abundant in maternal samples were more likely to be vertically transmitted **(Fig. S10)**. Taken together, these data suggest that vertical transmission may propagate lifestyle-dependent differences in microbiome composition (*33*).

Together our data show that infants from all lifestyles begin life with similar (*Bifidobacteria*-dominated) gut microbiota compositions, but subtle differences detected in early life compound over time. The minor taxa found by amplicon analysis to differentiate lifestyles at 0-6 months of life (*Bacteroides* in industrialized infants and *Prevotella* in non-industrialized infants) were the same taxa revealed by detailed metagenomic analysis to be the most commonly vertically transmitted. These data suggest that vertical transmission may be a mechanism by which microbiota change is propagated over generations in response to altered lifestyles (*34, 35*). Important differences were also discovered in the species composition and HMO-degradation genes of the initially-dominant *Bifidobacterium* communities, and recent studies of these same genes suggest that their depletion in industrialized infants could have long-term negative immune consequences (*25*). Crucially, in almost all analyses performed in this study, infants living transitional lifestyles display intermediate phenotypes between those of industrialized and non-industrialized infants. While not conclusive, this is an important piece of evidence pointing to lifestyle as a possible causative factor in infant microbiome assembly. The Hadza-specific discoveries reported in this work (including the finding of increased non-dyad vertical transmission among members of the same bush camp, a social structure with no equivalent among industrialized communities) exemplify the importance of studying people outside of industrialized nations, and highlights the need for additional studies to provide equity in microbiome understanding across global societies. Our results also highlight the question of whether lifestyle specific differences in the gut microbiome’s developmental trajectory predispose populations to diseases common in the industrialized world, such as those driven by chronic inflammation (*36, 37*).

## Acknowledgements

We acknowledge the numerous people and organizations who provided logistical support and conducted sample collection in the USA, Tanzania, and Nepal, including Dorobo Safaris, the Human Food Project, John Changalucha, Alphaxard Manjurano, Maria Gloria Domiguez-Bello, Michelle St. Onge, Allison Weakly and Yoshina Gautam. We thank David Relman and Chris Damman for helpful discussion and input throughout the project conceptualization and analysis. The content is solely the responsibility of the authors and does not necessarily represent the official views of the National Institutes of Health. This research utilizes data obtained by the TEDDY study group, a collaborative clinical study sponsored by the National Institute of Diabetes and Digestive and Kidney Diseases (NIDDK), National Institute of Allergy and Infectious Diseases (NIAID), National Institute of Child Health and Human Development (NICHD), National Institute of Environmental Health Sciences (NIEHS), and Centers for Disease Control and Prevention (CDC), and JDRF. The data from the TEDDY study reported here were supplied by the database of Genotypes and Phenotypes (dbGaP; Study Accession: phs001443.v1.p1), which is maintained by the National Center for Biotechnology Information (NCBI). This manuscript was not prepared in collaboration with investigators of the TEDDY study and does not necessarily reflect the opinions or views of the TEDDY study, dbGaP, or the NIDDK.

## Funding

This work was funded by grants from the National Institutes of Health (DP1-AT009892 and R01-DK085025 to JLS), an NSF Graduate Research Fellowship to DD (DGE-1656518) and to BDM (DGE-114747), a Stanford Graduate Smith Fellowship to DD, and National Institutes of Health grant F32DK128865 to MRO. JLS is a Chan-Zuckerberg Biohub Investigator. Research reported in this publication was supported by the National Institute Of Diabetes And Digestive And Kidney Diseases of the National Institutes of Health under Award Number F32DK128865. This project was supported by a grant from the Bill and Melinda Gates Foundation.

## Author contributions

Conceptualization: DD, ARJ, JLS

Genomic Sequencing: NN, BY, BDM, ST, DD

Methodology: DD, ARJ, MRO, MMC, BDM, SJ, SH, HW

Data analysis: DD, MRO, MMC, BDM, ST, SJ

Funding Acquisition: DD, JLS, EDS, ARJ, MRO

Supervision: EDS, JLS, ARJ, SH

Writing - original draft: DD, MRO, EDS, JLS

Writing - reviewing and editing: MRO, DD, ARJ, EDS, JLS, MMC

## Competing interests

The authors declare no conflict of interest.

## Data and materials availability

The authors declare that the data supporting the findings of this study are available within the paper and its supplementary information files. Metagenomic reads and genomes generated in this study are available under BioProject PRJEB27517. Accession numbers for individual samples and genomes are available in **Tables S2 and S7**.

## Supplementary Materials

Materials and Methods

Figures S1-S10

Tables S1-S7

References (38-71)

## Notes

### Competing Interest Statement

The authors have declared no competing interest.

