## Supplemental_Methods_Figures for "Robust Variation in Infant Gut Microbiome Assembly Across a Spectrum of Lifestyles"

**This PDF file includes:**

Materials and Methods

Figs S1 to S10

**Other Supplementary Materials for this manuscript include the following:**

Table S1 to S7

### Materials and Methods

#### Hadza sample gathering

Fecal samples from Tanzania were gathered in 2013-2014 as described previously by Smits et al. (*1*) and Fragiadakis et al. (*2*) after having described the study's intent and scope to and receiving consent from the Hadza participants; mothers provided consent on behalf of infants. A material transfer agreement with the National Institute for Medical Research in Tanzania ensures that stool samples collected are used solely for academic purposes. Permission for the study was obtained from the National Institute of Medical Research (MR/53i 100/83, NIMR/HQ/R.8a/Vol.IX/1542) and the Tanzania Commission for Science and Technology.

#### 16S rRNA Amplicon Processing

16S rRNA and metagenomic samples used in meta-analyses were collected from accession sources provided in their respective publications (SRA/dbGaP/mgRast/ENA sources available in **Table S1**). For amplicon sequence variant level analysis, samples were only included if the V4 region (515f,805r) was sequenced. For a study that used the V3-V4 region (341f, 805r) (*12*), bbduk was used to trim the reads to V4. The age for Nigeria (Bassa) infant samples is unreported (*12*), but correspondence with the authors of the study revealed that infants were approximately aged 2.5 years old. An age of 30 months was used for plotting the PCA analysis **(Fig. 1A)**, but due to the inability to assign a precise age these samples were excluded from all statistical analyses (including those based on the PCA plot).

To accurately estimate sequencing errors and allow for robust cross-study ASV comparisons, the merged paired-end DADA2 pipeline (*38*) was run on each sequencing run, from each study, separately with no trimLeft parameters. In cases where multiple samples were available from the same subject, a single sample from each sample was chosen to ensure that similar age distributions were present for all lifestyles **(Fig. S2)**. DADA2 (*38*) was used with default parameters to filter reads (rm.phix = TRUE), learn amplicon error rates, dereplicate amplicons, infer amplicon sequence variants (ASVs), merge paired reads, construct a sequence table, and remove chimeras. ASVs were assigned taxonomy against the DADA2 formatted GTDB 16Sr RNA database (01/16/2019 version; DOI 10.5281/zenodo.2541239).

Metagenomic library preparation and sequencing

Shotgun metagenome sequencing was performed on DNA extracted using the MoBio PowerSoil kit or phenol:chloroform:isoamyl alcohol as previously described (*1*, *2*). Technical replicates were included to assess differences between library preparation methods and sequencing platforms.

Libraries were prepared using half-reactions of the Nextera Flex kit, using a minimum of 10 ng of DNA as input and only 6 or 8 PCR cycles depending on input concentration to minimize PCR amplification bias. During PCR, a different 12 base pair unique dual-indexed barcode was added to each sample. Libraries were quantified using an Agilent Fragment Analyzer, and then size-selected using AMPure XP beads (Beckman), targeting a fragment length of 450bp (insert size of 350 bp).

Paired-end sequencing (2x140bp) was performed on a NovaSeq 6000 using S4 flow cells. Samples were randomized across sequencing runs and the same library was sequenced several times until the target depth was reached. The average sequencing depth for Hadza infant samples is 55.4 million paired-end reads (14.8 giga base-pairs).

Lifestyle classification

Samples were primarily classified by lifestyle using the U.N. Human Development Index, as previously reported (*15*, *39*). Industrialized infants are from urban industrialized populations in countries in the top 50% of the U.N. Human Development Index (HDI > 0.75) (*4*, *10*, *11*, *40*). Non-industrialized infants are from indigenous communities that are isolated, practice non-industrial means of farming (e.g., swidden), foraging, and hunting, and use rivers as a main water source (*12*, *14*). Transitional populations are from populations in the bottom 50% of the U.N. HDI and are not "non-industrialized" (*10–13*, *41*, *42*). One exception is the Karelia cohort, which, consistent with a previous report (*11*), is labeled as transitional since despite being within Russia it is an isolated autonomous republic with high rates of self-sufficient farming and relatively low sanitation practices compared to urbanized populations (*11*). The other exception is the cohort from Loreto, Peru, which was labeled as transitional given its location in the Amazon Rainforest and its previous classification as a "periurban" area (*10*). When available, breastfeeding practices of industrialized infants were assigned one of the four categories laid out by Stewart et. al. based on metadata provided by original publications (*4*).

Metagenome quality control and assembly

Public metagenomic samples used in meta-analysis all exceeded 1Gbp of total sequencing, and the same quality preprocessing steps, including removal of hg19 mapping reads, were applied to all samples. See **Table S1** and **Table S2** for accession numbers for public metagenomes used in this study.

Raw sequencing reads were demultiplexed and data from all sequencing runs for each sample were concatenated prior to analysis. Raw reads were processed using software in the BBtools suite (*43*). First, exact duplicate reads (subs=0) were marked using clumpify. Adapters and low-quality bases were trimmed using bbduk (trimq=16 minlen=55). Trimmed reads were mapped using BBmap against the human genome (hg19) with masks over regions conserved broadly in eukaryotes. Duplicate reads were removed. FastQC (*44*) was used to ensure processed reads were of sufficient quality with no modules reporting errors. BBMerge was used to merge reads that could be joined unambiguously using the recommended optimal settings (rem k=62 extend2=50 ecct vstrict) (*45*).

Metagenomes were assembled individually using metaSPAdes (*46*) (3.13) using unmerged forward/reverse reads as well as merged reads (-k 21,33,55,77) with error-correction enabled. QUAST (*47*) v5.0 was used to evaluate assembly size and contig metrics. Assemblies were filtered to contigs ≥ 1500 base-pairs for all subsequent analyses.

Unsupervised clustering

Principal Coordinates Analysis (PCoA) was used to perform unsupervised clustering of samples. Weighted PCoA was performed using the `dudi.pco` function in `ade4 (1.7-15)` in R, as outlined in Holmes & Huber Chapter on Multivariate Analyses (*48*), in order to correct for non-uniform distributions of infant sample age and cohort. With 16S rRNA data, ASVs were transformed with a log+1 transformation and weighted PCoA was performed on an unweighted UniFrac dissimilarity matrix. For metagenomic functional data, read counts were input as reads-perkilobase-million (RPKM), and weighted PCoA was performed on a Bray-Curtis dissimilarity matrix. Multivariate analysis was then only performed on genes in the top 25% of variance across the entire dataset (2,775,626/11,884,784 genes). For these functions the `ade4 (1.7-15)`, `vegan (2.5-6)`, and `phyloseq (1.32.0)` packages along with their dependencies were used (*49–51*).

<u>EnvFit analysis to determine significant covariates in multivariate analyses</u>

As previously described for assessing the effect of covariates on infant microbiota ordinations (*4*), the effect size and significance of each covariate was determined using the 'envfit' function in 'vegan'. For 16S rRNA, EnvFit used ordination performed using PCoA based on unweighted UniFrac dissimilarity. For metagenomic data, EnvFit used ordination performed using PCoA based on Bray-Curtis dissimilarity matrix. EnvFit significance was determined on 10,000 permutations.

<u>Microbial co-abundance groups (CAG) identification</u>

FastSpar was used to infer a correlation network on ASVs and calculate p-values (*17*), parameters included 1000 bootstrap correlations (--number 1000) and 1000 permutations for p-value estimates (--permutations 1000). ASVs were first agglomerated at a cophenetic distance of 0.05. Only correlations ≤0.2 or ≥0.2 with permuted p-values ≤0.05 were considered significant. The gap-statistic (*52*) was calculated using fviz_nclust in `factoextra (v1.0.7) to estimate the optimal number of clusters in the FastSpar inferred significant correlation matrix, which are designated as co-abundance groups (CAGs).

<u>Metagenomic functional analysis</u>

Single-sample assemblies were performed using metaSPAdes (v3.13.1). Open-reading frames (ORFs) were predicted using Prodigal (v2.6.3), using the metagenomic flag, resulting in 36,183,563 million complete genes. These ORFs were clustered into gene clusters using CD-Hit-EST (v4.8.1) at a 99% threshold nucleotide identity, where the shorter sequences were required to be at least 90% the length and 99% sequence identity to the longest sequence in the cluster, resulting in 11.98 million representative genes for this dataset. The abundance of each of these gene clusters was determined for each sample by aligning reads, using Bowtie2 (v2.3.5), from shotgun sequencing onto these representative ORF sequences. The gene abundance was analyzed as described in "EnvFit analysis to determine significant covariates in multivariate analyses", above.

Genome recovery and refinement

In order to assess the relatedness between all Hadza infant metagenomes, MASH sketches (-s 1000000 -k 32 -m 2) (*53*) were created from all reads in each metagenome, and sketches were compared in a pairwise manner. For each metagenomic assembly, reads from that sample and the nine next closest related samples were mapped using Bowtie2 (*54*) (--very-sensitive -X 1000). MetaBAT2 (*55*) (v2.13, default settings), and the resultant mappings were used to generate genome bins using contig depth information for all 10 samples. Genome bin quality was assessed using CheckM (v1.1.2) (*56*).

Bins were refined using MAGpurify v1 ((*57*) (using weighted mode for gc_content, tetra_freq, and coverage). The database used by Nayfach et al. (*57*) for conspecific analysis was augmented by adding all genome bins from this study that were >=95% complete and <=5% contaminated. Contigs flagged for removal by any module were removed from bins. Rarely, a module suggested the removal of >25% of a bin's length, and in such cases that module was turned off. Refined genomes that were ≥50% complete and <10% contaminated according to CheckM were retained for further analysis, consistent with MIMAG standards (*58*).

Assessment of genome novelty

UHGG (*21*) was used to determine which MAGs recovered from Hadza infants represent novel species. This analysis only considered Hadza Infant MAGs which passed the quality thresholds used to establish UHGG (Completeness ≥50%, Contamination <10%, and [Completeness - (5x Contamination)] > 50. Forty-one of 786 MAGs assembled from Hadza infants pass the MIMAG criteria and fail the UHGG criteria. The 745 MAGs recovered from Hadza infants that passed both sets of filters were clustered with UHGG species representatives using dRep (*59*) with the command "--S_algorithm fastANI --multiround_primary_clustering --clusterAlg greedy -ms 10000 -pa 0.9 -sa 0.95 -nc 0.30 -cm larger". This command approximates species-level clustering (*60*). Hadza Infant MAGs in clusters without a UHGG species representative were considered novel species. The phylum to which MAGs belong **(Fig. S5A)** was determined using GTDB (*61*). For this plot the phyla "Firmicutes_A" and "Firmicutes_C" are both referred to as "Firmicutes". Hadza Infant MAGs were clustered with MAGs assembled from Hadza Adults (*20*) to determine how many Hadza samples contain each novel species **(Fig. S6B)**.

<u>Creating species-representative genome database</u>

Hadza Infant MAGs passing the MIMAG standards were next combined with genomes from another study (*20*) to create a non-redundant species-level genome database to map reads to. Genomes were de-replicated into species-level groups using dRep (*59*) with the command "--S_algorithm fastANI --multiround_primary_clustering --clusterAlg greedy -ms 10000 -pa 0.9 -sa 0.95 -nc 0.30 -cm larger". This command assigns genomes that share ≥ 95% average nucleotide identity (ANI) over 30% of their length to the same species (*60*). MASH (-s 100) was then used to estimate the ANI between all genomes within each species-level group, and each genome was assigned a "centrality" score according to its average ANI to all other genomes in the group. The genome with the highest score was chosen as the representative for each species-level group according to the formula: score = (1*completeness) - (5*contamination) + (0.5*log10(ctg_N50)) + (1*log10(contig_bp)) + (2*(centrality-0.95)*100).

Centrality was calculated in the same way between all genomes in the UHGG genome database (v1.0) using the species-grouping described above (*21*). Species representatives were chosen using the same formula as above. Representatives from *de novo* genomes generated in this study and those from the UHGG database (v1.0) were next compared using dRep (--S_algorithm fastANI --multiround_primary_clustering --clusterAlg greedy -ms 10000 -pa 0.9 -sa 0.95 -nc 0.30 -cm larger). The genome with the highest score was chosen as the representative for each species-level group according to the formula: score = (1*completeness) - (5*contamination) + (0.5*log10(ctg_N50)) + (1*log10(contig_bp)). To account for edge-cases, chosen representatives were compared again using the same dRep command, and winners were again chosen using the same scoring criteria. Species-level group membership was back-propagated to the original bins. Taxonomy was determined for all species-level representative genomes using GTDB (v95) (*61*). The taxonomy reported by GTDB was used to establish the taxonomy of each MAG.

A single .fasta file was created by concatenating all species representative genomes. The UHGG *Bifidobacterium infantis* genome (GUT_GENOME095938) was replaced with genomes for *Bifidobacterium longum* subsp. *infantis* (*B. infantis*; accession AP010889.1) and *Bifidobacterium longum* subsp. *longum* (*B. longum*; accession AP010888.1) in order to measure these important subspecies separately. *Bifidobacterium longum* and *Bifidobacterium infantis* have historically been referred to as different subspecies of the same species (*62*), but more modern whole-genome based classification procedures have categorized them as separate species (*61*). In this study we refer to them as distinct species, consistent with the GTDB-based taxonomic classification used throughout this study.

InStrain relative abundance calculations

Reads from each sample were mapped to the concatenated file using Bowtie 2 with default settings. 'inStrain profile' and `inStrain quick_profile` was run on all resulting mapping files with inStrain v1.4.0 on default settings, the later of which was run with coverM version 0.6.0 (https://github.com/wwood/CoverM) (*28*). Detection of a species in a sample was defined as that species representative genome being present with ≥0.5 breadth (meaning that at least half of the bases in the genome were covered by at least a read). Genome relative abundance was calculated as ((genome length * genome coverage) / total metagenomic read base pairs). Results from `inStrain quick_profile` were used for all analyses unless otherwise noted.

VANISH / BloSSUM species classification

Species prevalence was first calculated by lifestyle within age bins. A species was defined as present if it was detected with ≥ 0.5 breadth. For each species with a prevalence of 30% in at least one lifestyle, a Fisher's exact test was performed on the prevalence of each species within age bins in the Hadza infants and industrialized infants. P values were corrected using Benjamini/Hochberg correction. Species significantly more prevalent in the Hadza than industrialized infants at any age bin were considered "VANISH" species, and species significantly more prevalent in industrialized infants than the Hadza at any age bin were considered "BloSSUM" species.

Mapping-based HMO analysis

Mapping-based HMO analysis **(Table S5**, **Fig. 2E)** was performed by mapping metagenomic samples against the *Bifidobacterium longum* subsp. *infantis* ATCC 15697 genome (*B. infantis;* accession CP001095.1) using Bowtie2. To identify HMO-degradation genes, a table linking gene activity to genomic location was retrieved from a previous publication (*25*) (filename "humann2_HMO_annotation.csv"). Open reading frames (ORFs) were identified on the *B. infantis* ATCC 15697 genome using Prodigal (*63*), and Diamond (*64*) was used to match Prodigal's ORFs to those listed in "humann2_HMO_annotation.csv" **(Table S5)**. Mappings were profiled using InStrain (*28*). Genes with breadth greater than 0.4 and coverage greater than half the number of giga base-pairs in the metagenome were considered present. The later threshold was chosen to ensure that samples with more sequencing depth do not have higher rates of gene detection, and was chosen as it is the minimum coverage needed for the most shallowly-sequenced sample (1 Gbp) to detect a gene at 0.4 breadth (*28*). HMO clusters were considered present if all genes in the HMO degradation cluster were present. Prevalence was calculated as the percentage of metagenomes in a given lifestyle and age range with a HMO degradation cluster present.

Bifidobacterium species isolation and sequencing

Seven Hadza infant fecals samples were chosen for isolation of Bifidobacterium (samples 2487, 2261, 2614, 1834, 1932, 1947, and 1936). Samples were plated on BSM Agar (Millipore Sigma SKU 88517-250G-F) and custom Bifidobacterium minimal media under anaerobic conditions. Custom media was based on DSMZ medium 104 (104. PYG MEDIUM, © 2020 DSMZ GmbH) and previously published media (*65*) with 2′-Fucosyllactose as the sole carbon source. Strains were identified via high-throughput MALDI-TOF mass spectrometry of whole colonies, and subsequent matching of spectra to a reference library with a MALDI Biotyper System (Bruker), following manufacturer's instructions and including formic acid lysis. Many isolated strains are not currently amenable to freezer storage and liquid-culture-based propagation in isolation. Colonies identified as Bifidobacterium were subjected to DNA sequencing. Genomic DNA was extracted from pure cultures using a DNeasy UltraClean 96 Microbial Kit (Qiagen 10196-4) or DNeasy Blood and Tissue Kit (Qiagen 69504) kit. Libraries were prepared using half-reactions of the Nextera Flex kit, libraries were quantified using an Agilent Fragment Analyzer, and then size-selected using AMPure XP beads (Beckman), targeting a fragment length of 450bp (insert size of 350 bp). Reads were processed and genomes were assembled as described above. GTDB (release 202) was used for species-level taxonomic assignment, and only species identified as *Bifidobacterium infantis* were used in this analysis.

Genome-based Bifidobacterium analysis

Public *Bifidobacterium infantis* isolates were identified by first downloading all Bifidobacterium genomes on NCBI as of July 12, 2021 using the program "ncbi-genome-download" (https://github.com/kblin/ncbi-genome-download). For all genomes of the "Chromosome" and "Complete Genome" assembly level, GTDB release 202 was run to establish current species-level taxonomy. Twelve genomes were identified as *Bifidobacterium infantis*, and these twelve genomes are enumerated in **Table S7**. Public *Bifidobacterium* MAGs were identified in the UHGG genome database (*21*). Only genomes with Contamination ≤ 5% and Completeness ≥ 80% were considered. GTDB release 202 was run on all genomes satisfying these chritrea and classified as Bifidobacterium in UHGG, and this information was used for species-level taxonomic assignments. ORFs were identified in all genomes using Prodigal (*63*).

HMO genes were identified using the program DIAMOND (*64*) to compare gene amino acid sequences to the HMO degradation genes present in *Bifidobacterium infantis* ATCC 15697 (HMO gene identification described above). Genes with percentID ≥ 50 and e-value ≤ 0.0001 were considered to be the same. Genomes with all genes present in an HMO-degradation cluster were considered to encode that cluster.

CAZymes were identified in genomes using the program hmmscan (*66*) to compare gene amino acid sequences against dbCAN2 HMMs (V9; filename dbCAN-HMMdb-V9.txt). The dbCAN2 script "hmmscan-parser.sh" was run on resultant files, and custom code was used to parse CAZyzme family names from HMM names.

Pfams were identified in genomes using the program hmmsearch (*66*) to compare gene amino acid sequences against Pfam HMMs (v32.0; filename "Pfam-A.hmm"). The gathering cutoff was used to filter results, and the program “cath-resolve-hits.ubuntu14.04” (*67*) was used to collapse the domain matches down to the best, non-overlapping subset.

Gene clusters were identified in genomes via *de novo* clustering using MMseqs2 (*68*). All protein sequences from all *Bifidobacterium infantis* genomes analyzed in this study were compared using the MMseqs2 easy-linclust workflow with the additional arguments “--cov-mode 1 -c 0.8 --kmer-per-seq 80 --min-seq-id 0.95”. The resultant files were parsed using custom scripts to identify clusters with differential prevalence in genomes from different lifestyles.

Phylogenetic analysis was performed using the program GToTree v1.5.36 (*69*). Hadza isolate genomes, all MAGs identified as *Bifidobacterium infantis*, and public reference genomes *Bifidobacterium longum* strain 35624 (BioSample SAMN04254466) and *Bifidobacterium longum* subsp. *infantis* ATCC 15697 (*B. infantis*; BioSample SAMD00060952) were included as input. Default parameters and bacterial single copy genes were used (-H Bacteria). The phylogenetic tree was visualized using iTol (*70*).

<u>InStrain strain tracking analysis</u>

For InStrain strain tracking analysis (*28*), genome detection was defined as having minCov breadth ≥0.5 (meaning that at least half of the bases in the genome were covered by at least 5 reads) as measured using `inStrain profile`. Each species detected in more than one individual was compared using ‘inStrain compare’. A distance matrix was created for each species based on popANI values, and this matrix was used to cluster subspecies into a number of individual strains using ‘average’ hierarchical clustering with a threshold of 99.999% popANI using the Scipy cluster package. The number of strains shared between sample pairs was calculated based on this strain definition. The percentage of shared strains shared between dyads was calculated as the number of shared strains divided by the number of strains found in the infant sample..

To perform a comparison of vertically transmitted strains that is robust to differences in sequencing depth, all infant and maternal samples were subset *in silico* to a total of 4 Gbp using seqtk (*71*). Samples with less than 4 Gbp of sequencing were excluded from this analysis. The number of read pairs retained for each sample was calculated as (4,000,000,000 / average read pair length) as enumerated in **Table S2**. Rarefied sets of paired reads were then analyzed in the same way as unrarefied reads, as described above (reads were mapped to the same set of species representative genomes, inStrain profile was run using the same parameters, and strains were identified using the same workflow).

To identify taxa that are significantly enriched among strains in the Hadza / Swedish populations that are shared among dyads **(Fig. 3C)**, a Fisher's exact test was run on the table [[number of vertically transmitted strains of this taxa in Swedish infants, number of non-vertically transmitted strains of this taxa in Swedish infants], [number of vertically transmitted strains of this taxa in Hadza infants, number of non-vertically transmitted strains of this taxa in Hadza infants]] using the Scipy python package. The test was run on the top 3 most abundant phylum-level and genus-level taxa, and P values were corrected using Benjamini/Hochberg correction. For this plot the phyla "Firmicutes_A" and "Firmicutes_C" are both referred to as "Firmicutes".

Supplemental Notes and References

38. B. J. Callahan, P. J. McMurdie, M. J. Rosen, A. W. Han, A. J. A. Johnson, S. P. Holmes, DADA2: High-resolution sample inference from Illumina amplicon data. *Nat. Methods*. **13**, 581–583 (2016).

39. Human Development Data Center, (available at http://www.hdr.undp.org/en/data).

40. F. Bäckhed, J. Roswall, Y. Peng, Q. Feng, H. Jia, P. Kovatcheva-Datchary, Y. Li, Y. Xia, H. Xie, H. Zhong, M. T. Khan, J. Zhang, J. Li, L. Xiao, J. Al-Aama, D. Zhang, Y. S. Lee, D. Kotowska, C. Colding, V. Tremaroli, Y. Yin, S. Bergman, X. Xu, L. Madsen, K. Kristiansen, J. Dahlgren, J. Wang, Dynamics and Stabilization of the Human Gut Microbiome during the First Year of Life. *Cell Host Microbe*. **17**, 852 (2015).

41. A. Tett, K. D. Huang, F. Asnicar, H. Fehlner-Peach, E. Pasolli, N. Karcher, F. Armanini, P. Manghi, K. Bonham, M. Zolfo, F. De Filippis, C. Magnabosco, R. Bonneau, J. Lusingu, J. Amuasi, K. Reinhard, T. Rattei, F. Boulund, L. Engstrand, A. Zink, M. C.

Collado, D. R. Littman, D. Eibach, D. Ercolini, O. Rota-Stabelli, C. Huttenhower, F. Maixner, N. Segata, The Prevotella copri Complex Comprises Four Distinct Clades Underrepresented in Westernized Populations. *Cell Host Microbe*. **26**, 666–679.e7 (2019).

42. E. C. Pehrsson, P. Tsukayama, S. Patel, M. Mejía-Bautista, G. Sosa-Soto, K. M. Navarrete, M. Calderon, L. Cabrera, W. Hoyos-Arango, M. T. Bertoli, D. E. Berg, R. H. Gilman, G. Dantas, Interconnected microbiomes and resistomes in low-income human habitats. *Nature*. **533**, 212–216 (2016).

43. B. Bushnell, BBTools software package. *URL http://sourceforge. net/projects/bbmap*. **578**, 579 (2014).

44. S. Andrews, Others, FastQC: a quality control tool for high throughput sequence data (2010).

58. R. M. Bowers, N. C. Kyrpides, R. Stepanauskas, M. Harmon-Smith, D. Doud, T. B. K. Reddy, F. Schulz, J. Jarett, A. R. Rivers, E. A. Eloe-Fadrosh, S. G. Tringe, N. N. Ivanova, A. Copeland, A. Clum, E. D. Becraft, R. R. Malmstrom, B. Birren, M. Podar, P. Bork, G. M. Weinstock, G. M. Garrity, J. A. Dodsworth, S. Yooseph, G. Sutton, F. O. Glöckner, J. A. Gilbert, W. C. Nelson, S. J. Hallam, S. P. Jungbluth, T. J. G. Ettema, S. Tighe, K. T. Konstantinidis, W.-T. Liu, B. J. Baker, T. Rattei, J. A. Eisen, B. Hedlund, K. D. McMahon, N. Fierer, R. Knight, R. Finn, G. Cochrane, I. Karsch-Mizrachi, G. W. Tyson, C. Rinke, Genome Standards Consortium, A. Lapidus, F. Meyer, P. Yilmaz, D. H. Parks, A. M. Eren, L. Schriml, J. F. Banfield, P. Hugenholtz, T. Woyke, Minimum information about a single amplified genome (MISAG) and a metagenome-assembled genome (MIMAG) of bacteria and archaea. *Nat. Biotechnol.* **35**, 725–731 (2017).

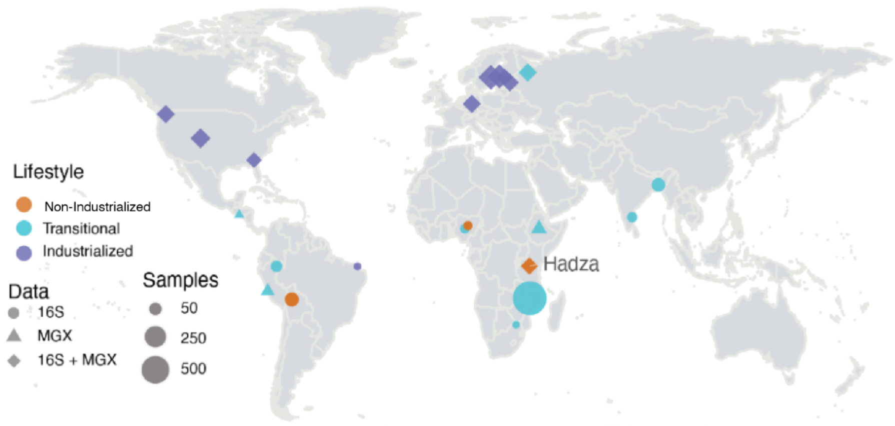


**16S rRNA**

| | 0-6 | 6-12 | 12-24 | 24-36 |
|---|---|---|---|---|
| Industrialized | 156 | 284 | 274 | 181 |
| Non-Industrialized | 28 | 31 | 26 | 25 |
| Transitional | 205 | 202 | 239 | 240 |

| | 0-6 | 6-12 | 12-24 | 24-36 |
|---|---|---|---|---|
| Bangladesh | 10 | 7 | 11 | 17 |
| Bolivia (Tsimane) | 16 | 22 | 10 | 0 |
| Brazil | 1 | 1 | 4 | 1 |
| Colorado (US) | 19 | 35 | 42 | 18 |
| Estonia | 12 | 12 | 18 | 20 |
| Finland | 52 | 85 | 69 | 70 |
| Georgia (US) | 4 | 16 | 11 | 7 |
| Germany | 16 | 23 | 24 | 8 |
| India | 9 | 3 | 2 | 0 |
| Malawi | 174 | 181 | 181 | 179 |
| Nigeria (Bassa) | 0 | 0 | 0 | 0 |
| Nigeria (Urban) | 0 | 1 | 6 | 5 |
| Peru | 11 | 1 | 8 | 2 |
| Russia (Karelia) | 0 | 7 | 28 | 36 |
| South Africa | 1 | 2 | 3 | 1 |
| Sweden | 38 | 93 | 80 | 47 |
| Tanzania (Hadza) | 12 | 9 | 16 | 25 |
| Washington (US) | 14 | 19 | 26 | 10 |

**Metagenomics**

| | 0-6 | 6-12 | 12-24 | 24-36 |
|---|---|---|---|---|
| Industrialized | 151 | 134 | 181 | 103 |
| Non-Industrialized | 13 | 9 | 10 | 7 |
| Transitional | 14 | 15 | 35 | 10 |

| | 0-6 | 6-12 | 12-24 | 24-36 |
|---|---|---|---|---|
| Colorado (US) | 7 | 14 | 23 | 13 |
| El Salvador | 1 | 0 | 2 | 0 |
| Estonia | 7 | 10 | 16 | 9 |
| Ethiopia | 6 | 8 | 7 | 4 |
| Finland | 31 | 28 | 42 | 33 |
| Georgia (US) | 6 | 8 | 14 | 4 |
| Germany | 7 | 12 | 18 | 9 |
| Peru | 1 | 1 | 4 | 2 |
| Russia (Karelia) | 6 | 6 | 22 | 4 |
| Sweden | 81 | 51 | 50 | 24 |
| Tanzania (Hadza) | 13 | 9 | 10 | 7 |
| Washington (US) | 12 | 11 | 18 | 11 |

**Figure S1. Global map of infant samples used in this study**. Points are colored by lifestyle; shapes indicate data type; sizes indicate sample number used in this study. Cohort information and information including sample counts, subject counts, and references are available in Google My Map (https://bit.ly/3iq5r8v).

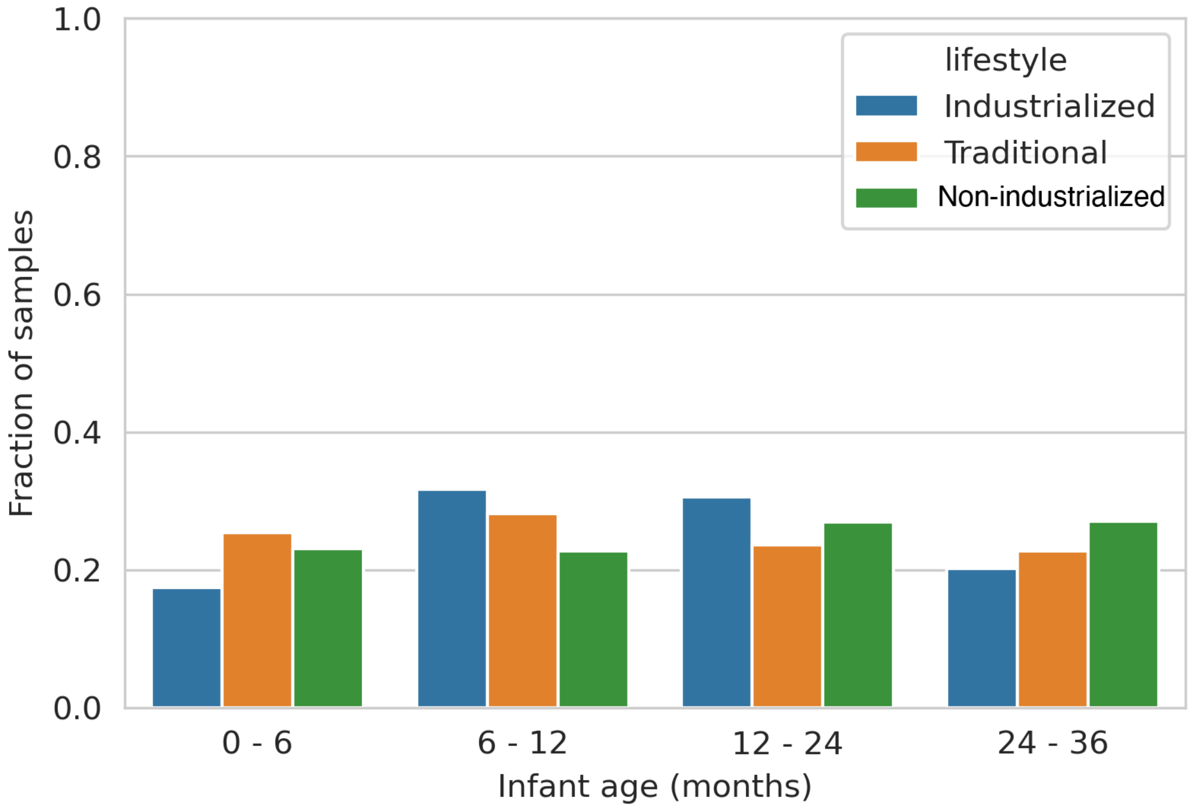


**Figure S2. Sample age proportion by lifestyle.** For each lifestyle, the fraction of infant fecal 16S samples in each age bin.

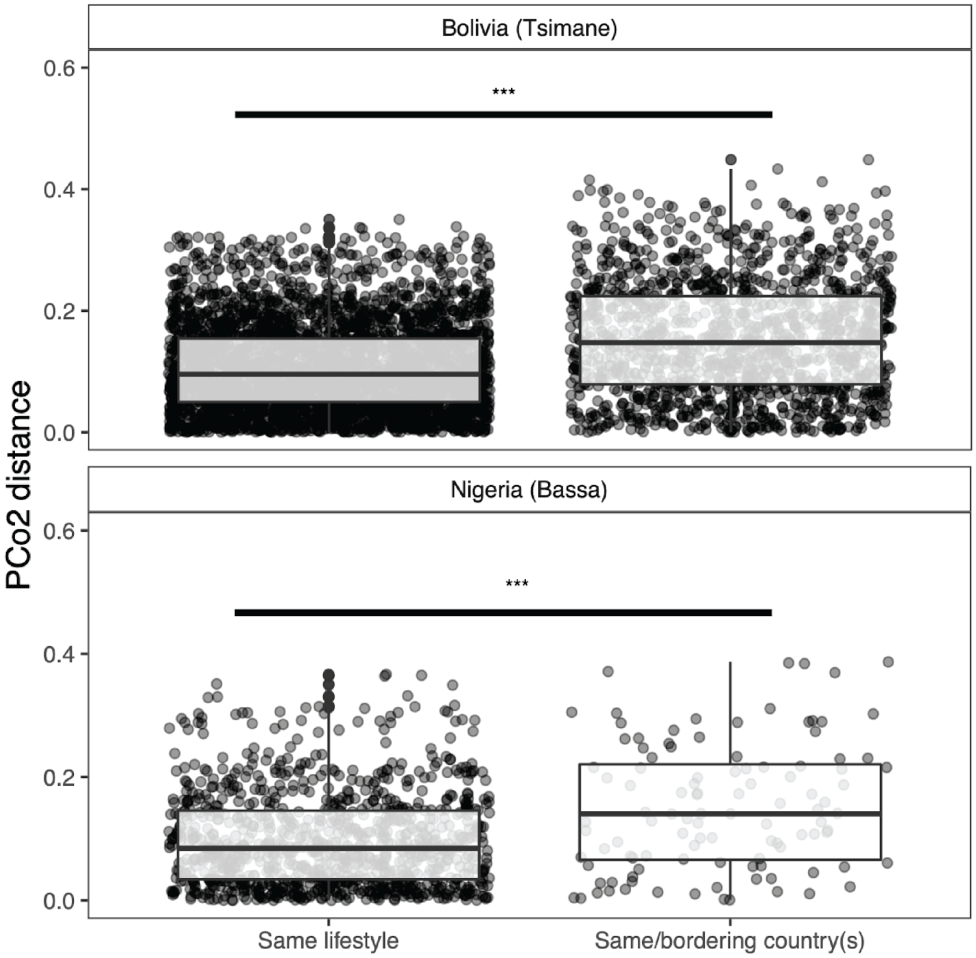


**Figure S3. Shared lifestyle affects microbiota composition more than geographic proximity.** Pairwise distances between samples on PCo2 from **Fig. 1A**; comparisons within the same lifestyle do not include samples from the same cohort. Bolivia (Tsimane, top panel) compared to the same lifestyle (Tanzania (Hadza) and Nigeria (Bassa); n=432) or to the same/bordering country with a different lifestyle (Brazil (industrialized) and Peru (transitional); n=1392) (Wilcoxon-rank test; P=2.2e-16; W=210364). Nigeria (Bassa, bottom panel) compared to the same lifestyle (Tanzania (Hadza) and Bolivia (Tsimane); n=432) or compared to the same/bordering country with different lifestyles (urban Nigeria population, industrialized; n=108) (Wilcox-rank test; P=0.0001; W=17696) (*** = P < 0.01).

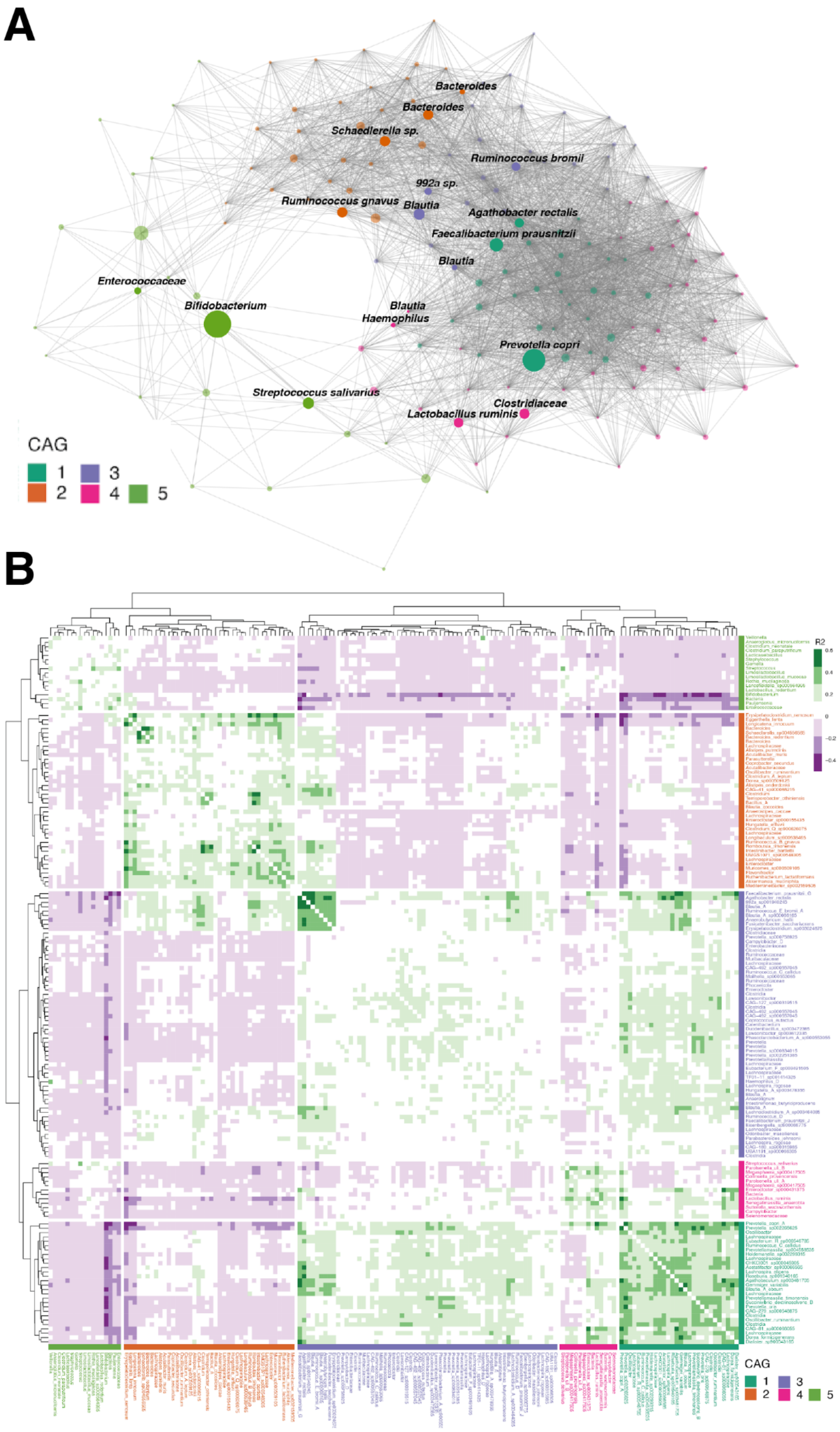
A
Bacteroides
Bacteroides
Schaedlerella sp.
Ruminococcus bromii
992a sp.
Ruminococcus gnavus
Blautia
Agathobacter rectalis
Faecalibacterium prausnitzii
Blautia
Enterococcaceae
Bifidobacterium
Blautia
Haemophilus
Prevotella copri
Streptococcus salivarius
Clostridiaceae
Lactobacillus ruminis
CAG
1
2
3
4
5
B
R2
0.6
0.4
0.2
0
−0.2
−0.4
CAG
1
2
3
4
5

**Figure S4. Correlation heatmap and network representation of CAGs. (A)** Gap-statistic computed co-abundance groups (CAGs) from $R^2$ correlation values across all 16S rRNA abundance across all 1,900 samples are colored by blocks. Cell values show $R^2$ correlations calculated with FastSpar (see methods). **(B)** Taxa within five CAGs. Size of nodes are average relative abundance in age group and lifestyle in which that taxon has the highest abundance. Edges show significant positive correlations with an $R^2$ greater than 0.1. Labeled taxa are those with the highest average within sample relative abundance by CAG.

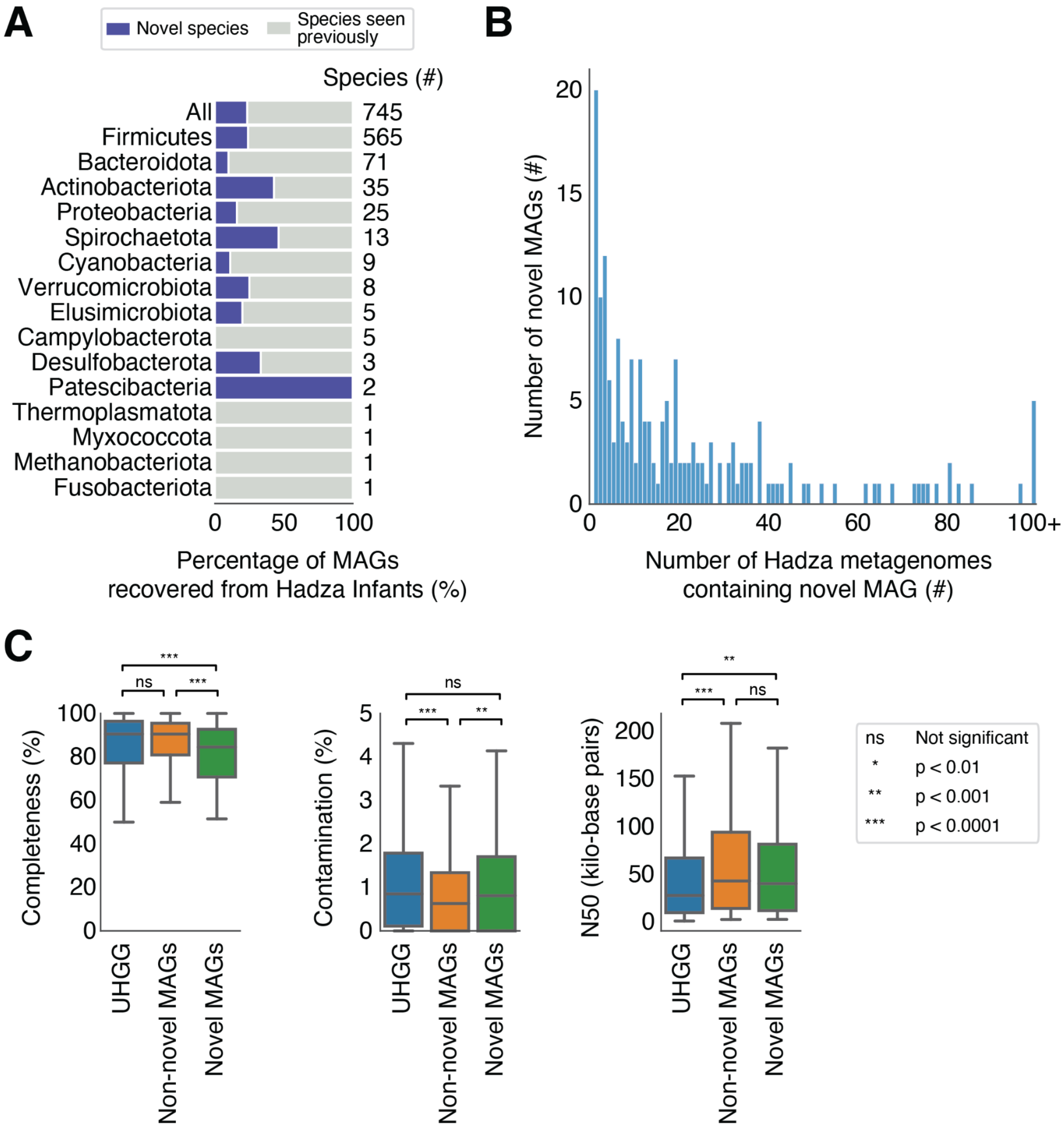


**Figure S5. MAGs recovered from the Hadza infant microbiome are phylogenetically diverse and high quality. (A)** Percentage of species assembled from Hadza infant samples, grouped by phylum, that are novel relative to UHGG. Numbers on the right indicate total (novel plus known) species by phylum. **(B)** Histogram of the number of Hadza gut metagenomes (infant and adult) containing each novel Hadza infant MAG. Of the 175 novel MAGs from Hadza infants, 155 are assembled from > 1 Hadza metagenome. **(C)** Comparison of genome quality metrics of all genomes in UHGG (n=286,997; "UHGG"), novel MAGs recovered from Hadza infants (n=175; "Novel MAGs"), and other MAGs recovered from Hadza infants (n=570; "Non-novel MAGs"). P-values from Wilcoxon rank-sum test. Effect sizes, reported as Cohen's d, for UHGG vs. Non-novel MAGs are -0.10, 0.27, and -0.08, for UHGG vs. Novel MAGs are 0.33, 0.01, and -0.04, and for Non-novel MAGs vs. Novel MAGs are 0.47, -0.28, and 0.08 for Completeness, Contamination, and N50, respectively.

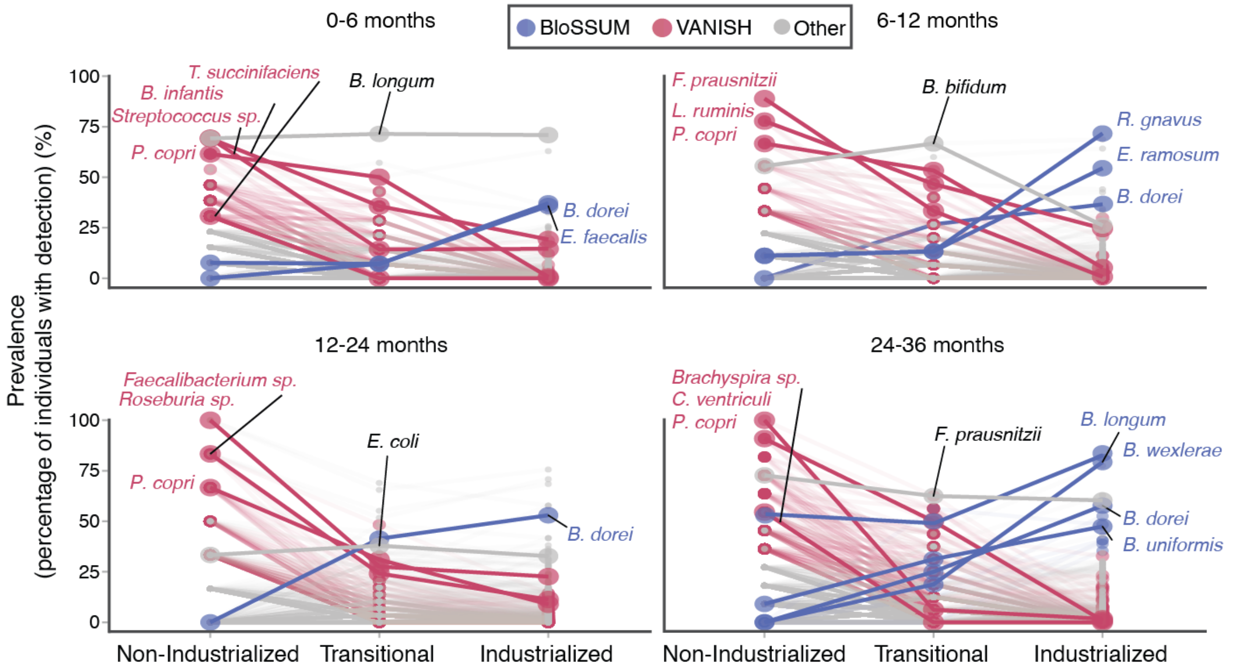


**Figure S6. VANISH and BloSSUM taxa across age windows.** Fractional prevalence of species across lifestyles, faceted by age group. Select VANISH (red) and BloSSUM (blue) (those with lowest adj-P) species are highlighted. "Other" (gray) taxa are those that are not significantly different by age and lifestyle.

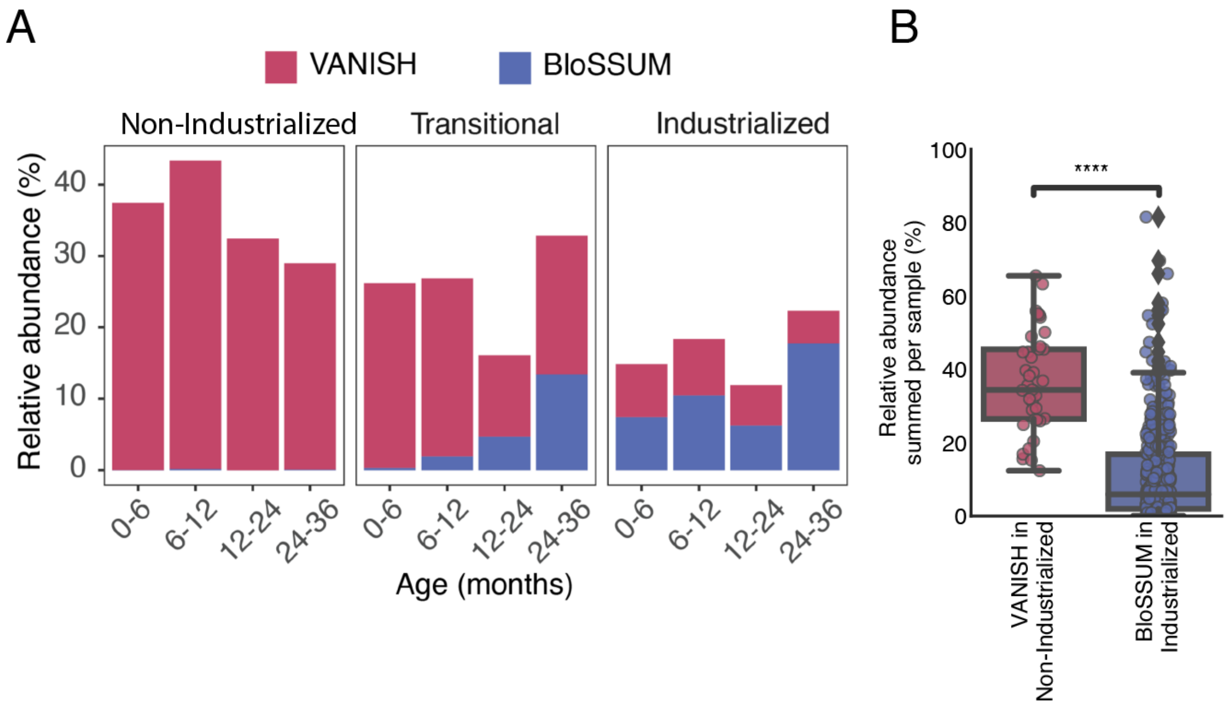


**Figure S7. Cumulative relative abundance of VANISH and BloSSUM taxa in infants by age and lifestyle. (A)** The average abundance of VANISH and BloSSUM taxa in infants from different lifestyles over time. **(B)** Over the first 3 years of life there is a significantly higher fraction of VANISH taxa in traditional infants then there are BloSSUM taxa in industrialized infants (P=8.2e-21; Mann-Whitney U statistic=1.7E3; n=569 industrialized infants; n=39 traditional infants; Wilcoxon rank-sum test).

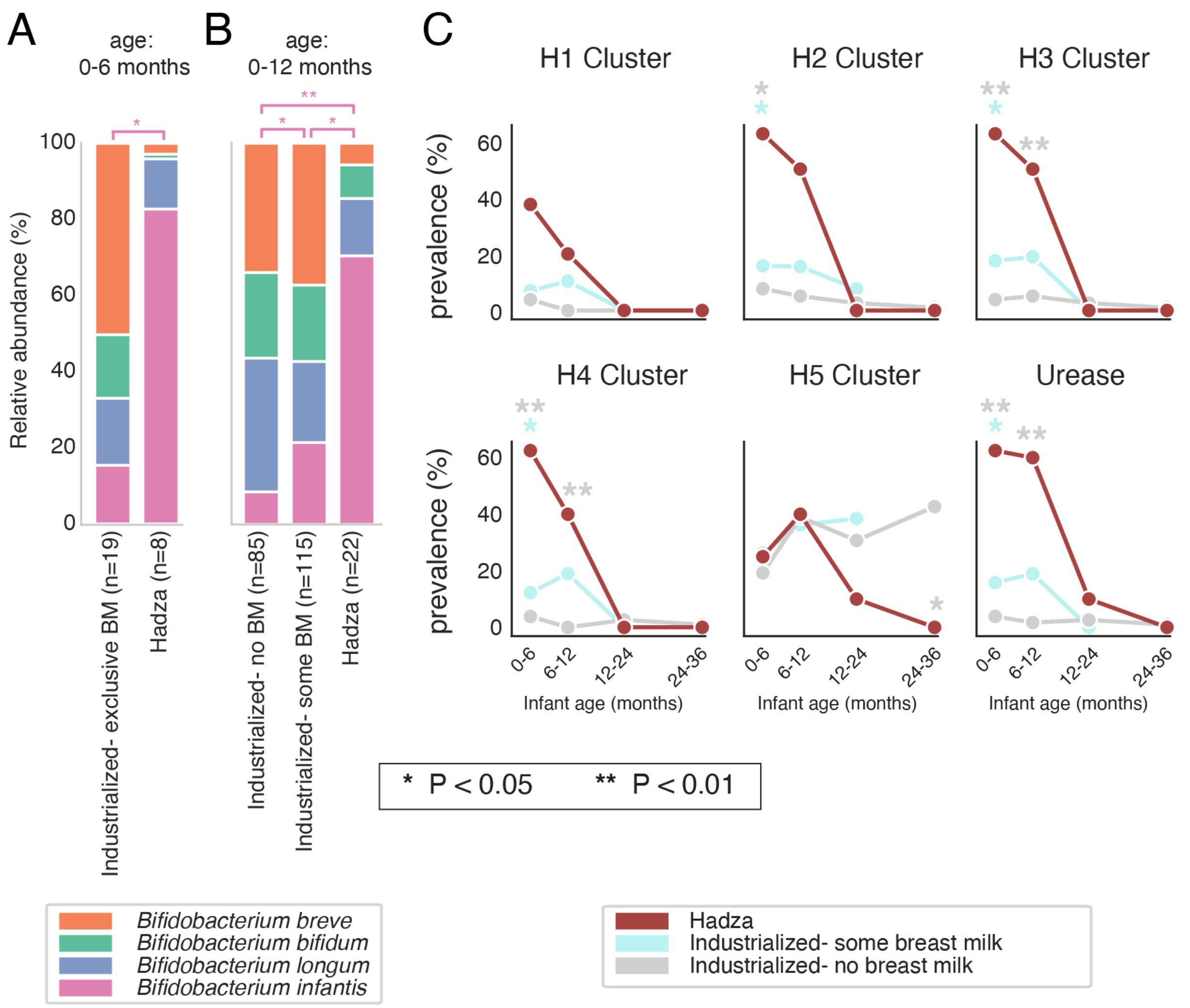


**Figure S8.** ***Bifidobacterium infantis*** **and HMO-degradation cassette prevalence among infants which do and do not consume breast milk. (A, B)** Relative representation of four common *Bifidobacterium* species in infants **(A)** ≤ 6 months old and **(B)** ≤ 1 year old. BM=breast milk. "Industrialized- no BM" includes industrialized infants that have never consumed breastmilk or no longer consume breast milk; "Industrialized- some breast milk" includes industrialized infants that exclusively or partially consume breast milk; "Industrialized- exclusive breast milk" includes industrialized infants that exclusively consume breast milk. Asterisks at top indicate statistical significance of *Bifidobacterium infantis* relative abundance between groups (Wilcoxen rank-sums test). (B) Prevalence of HMO-utilization clusters across the same groups as described in (B). Clusters are considered present if all genes in the cluster are detected above a variable coverage threshold. Blue asterisks indicate statistical significance of non-industrialized versus Industrialized- some breast milk, grey asterisks indicate statistical significance of Hadza versus Industrialized- no breast milk (Fisher’s exact test with false discovery rate correction).

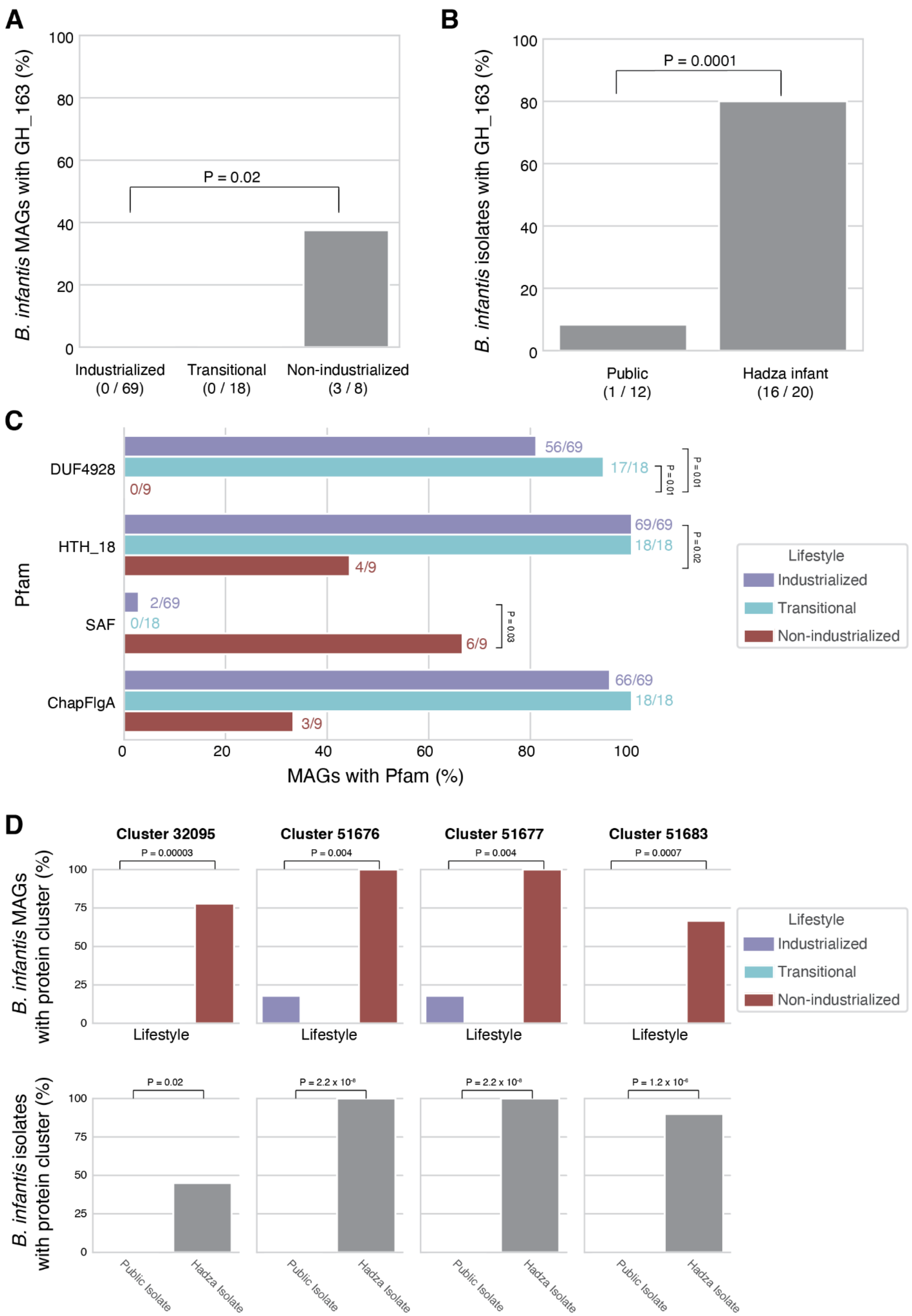


**Figure S9. *Bifidobacterium infantis* genomes exhibit lifestyle-associated differences.** The percentage of *B. infantis* MAGs **(A)** and *B. infantis* isolate genome sequences **(B)** which contain

CAZyme GH_163. MAGs are compared based on the lifestyle of the infant from which the MAG was assembled, and isolates are compared based on whether they were isolated from Hadza Infant stool samples or downloaded from the NCBI public genome database. The number of genomes with GH_163 and the total number of genomes are shown below the x-axis for each group. P values are from Fisher's exact test with false discovery rate correction for testing all CAZyme families present in at least one genome. Odds ratio for Industrialized vs. Non-industrialized in subpanel A=0; odds ratio for Public vs. Hadza infant in subpanel B=0.02. **(C)** The percentage of *B. infantis* MAGs that contain the Pfam domains listed on the y-axis. The number of genomes with each Pfam and the total number of genomes are shown to the right of the y-axis for each group. P values are from Fisher's exact test with false discovery rate correction for testing all Pfams. All Pfams with significant differences between industrialized and non-industrialized lifestyles are shown, as well as ChapFlgA. While not significant after false discovery rate correction, ChapFlgA is a similar family as SAF; both are involved in flagellar assembly. Odds ratios for industrialized vs. transitional are 0.25, NaN, inf., and 0; for industrialized vs. non-industrialized are inf., inf., 0.06, and 44, and for transitional vs. non-industrialized are inf., inf., 0, and inf. for CUF4928, HTH_18, SAF, and ChapFlgA, respectively. **(D)** The percentage of *B. infantis* MAGs (top row) and *B. infantis* isolate genome sequences (bottom row) that contain various *de novo* protein clusters (columns). P values are from Fisher's exact test with false discovery rate correction for testing all clusters. Odds ratio for all Industrialized vs. Non-industrialized (top) and all Public isolate vs. Hadza Isolate comparisons (bottom) = 0. All clusters with significance among both MAGs and isolates are shown. Representative proteins for the *de novo* clusters match the following Pfams: 32095 hits "Penicillin-binding Protein dimerisation domain" and "Penicillin binding protein transpeptidase domain"; 51676 and 51677 both hit "ABC transporter" and "ABC transporter transmembrane region; 51683 hits "Phage integrase, N-terminal SAM-like domain" and "Phage integrase family".

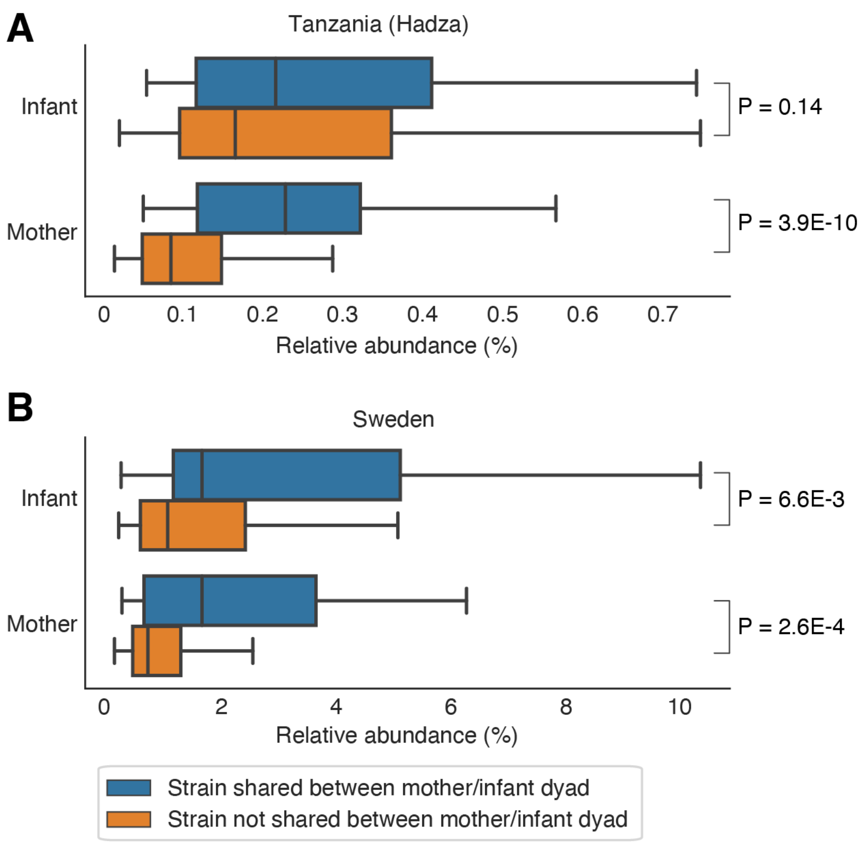


**Figure S10. More highly abundant strains are more likely to be shared between mother/infant dyads. (A, B)** The relative abundance of bacteria in the infant (top) and maternal (bottom) microbiomes that are shared (blue) and not shared (orange) between mother/infant dyads in the **(A)** Hadza and **(B)** Swedish populations within rarefied metagenomes. Among the Hadza, n=48 and 311 shared and not shared bacteria in the infant microbiome (P=0.14, O=1.48) and n=48 and 429 shared and not shared bacteria in the maternal microbiome (P=3.86E-10, O=6.3). Among Swedish dyads, n=32 and 376 shared and not shared bacteria in the infant microbiome (P=6.6E-3, O=2.7) and n=32 and 337 shared and not shared bacteria in the maternal microbiome (P=2.6E-4, O=3.6). P values and effect sizes from Wilcoxon-rank test.

### Supplemental tables

**Table S1**: Sample information and study accession numbers

**Table S2**: Metagenomic sample metadata

**Table S3**: Metagenomic sample abundance

**Table S4**: Prevalence and abundance of all taxa with VANISH/BloSSUM/Other status across age bins

**Table S5**: Information on HMO gene identification and HMO gene abundance

**Table S6**: Vertical transmission rates, both rarefied and non-rarefied

**Table S7:** Information on recovered genomes
